# Mechano-responsiveness of fibrillar adhesions on stiffness-gradient gels

**DOI:** 10.1101/775809

**Authors:** Nuria Barber-Pérez, Maria Georgiadou, Camilo Guzmán, Aleksi Isomursu, Hellyeh Hamidi, Johanna Ivaska

**Affiliations:** Turku Bioscience Centre, University of Turku and Åbo Akademi University, FIN-20520 Turku, Finland; Department of Biochemistry, University of Turku, FIN-20520 Turku, Finland

## Abstract

Fibrillar adhesions are important structural and adhesive components in fibroblasts that are critical for fibronectin fibrillogenesis. While nascent and focal adhesions are known to respond to mechanical cues, the mechanoresponsive nature of fibrillar adhesions remains unclear. Here, we used ratiometric analysis of paired adhesion components to determine an appropriate fibrillar adhesion marker. We found that active α5β1-integrin exhibits the most definitive fibrillar adhesion localisation compared to other proteins, such as tensin-1, reported to be in fibrillar adhesions. To elucidate the mechanoresponsiveness of fibrillar adhesions, we designed and fabricated thin polyacrylamide (PA) hydrogels, embedded with fluorescently labelled beads, with physiologically relevant stiffness gradients using a cost-effective and reproducible technique. We generated a correlation curve between bead density and hydrogel stiffness, thus allowing the use of bead density as a readout of stiffness, eliminating the need for specialised knowhow including atomic force microscopy (AFM). We find that stiffness promotes the growth of fibrillar adhesions in a tensin-dependent manner. Thus, the formation of these extracellular matrix-depositing structures is coupled to the mechanical parameters of the cell environment and may enable cells to fine-tune their matrix environment in response to alternating physical conditions.

## Introduction

It has been known for nearly two decades that cultured fibroblasts form distinct types of ECM adhesions, the short-lived peripheral nascent adhesions, which are superseded by actin-tethered focal adhesions, and lastly mature, centrally located, elongated fibrillar adhesions (Katz et al., 2000; Zamir et al., 1999). Fibrillar adhesions mediate fibronectin remodelling and the formation of fibrils, which guide the deposition of other matrix components such as collagens, fibrillin, fibulin and tenascin-C (Chung and Erickson, 1997; Dallas et al., 2005; Kadler et al., 2008; McDonald et al., 1982; Sabatier et al., 2009; Saunders and Schwarzbauer, 2019; Singh et al., 2010; Sottile and Hocking, 2002; Twal et al., 2001; Velling et al., 2002) and are thus important for the formation of the extracellular matrix (ECM). Fibrillar adhesions are partly defined by the presence of α5β1-integrin and tensin and the absence of other integrins heterodimers (Pankov et al., 2000; Zamir et al., 2000). Ligand-bound α5β1-integrin translocates centripetally out of focal adhesions along the actin cytoskeleton, organizing bound fibronectin into fibrils (Pankov et al., 2000; Zamir et al., 2000). Active (i.e. fully primed or ligand occupied) α5β1-integrin is recognized by the SNAKA51 antibody and co-localizes with fibronectin in fibrillar adhesions (Clark et al., 2005).

The assembly and dynamics of nascent and focal adhesions, and thus cellular functions such as cell migration, spreading and differentiation, are known to be regulated by both chemical and mechanical cues (e.g. viscoelastic properties, tensile forces) emanating from the ECM (Choi et al., 2012; Hadden et al., 2017; Hetmanski et al., 2019; Holle et al., 2016; Lo et al., 2000; Martino et al., 2018; Pelham and Wang, 1998; Wang et al., 2012). Although HIC-5, a paxillin family member, was recently shown to be required for the formation of tensin-1-positive fibrillar adhesions on rigid substrates (Goreczny et al., 2018), it still remains unclear whether fibrillar adhesions are also susceptible to changes in ECM elasticity.

Polyacrylamide(PA)-based hydrogels are the most commonly used *in vitro* cell culture platforms to study cellular behaviour in response to ECM elasticity, often referred to as stiffness or rigidity (Caliari and Burdick, 2016; Engler et al., 2006; Rowlands et al., 2008; Wen et al., 2014). These PA-hydrogels are usually generated with a uniform stiffness and while very informative for elucidating some of the molecular details regulating cell behaviour, are not representative of the *in vivo* situation. *In vivo*, the cellular microenvironment is extremely heterogeneous, not only in composition, but also in terms of stiffness (Young et al., 2016). Several different methods have been developed to generate stiffness gradient hydrogels that more closely mimic the mechanical heterogeneity observed *in vivo*, all with their own advantages and disadvantages (Chao et al., 2014; Hartman et al., 2016; Isenberg et al., 2009; Vincent et al., 2013). The main limitations include time-consuming, complex methodologies, or the need for specialised equipment not easily accessible in every laboratory. Moreover, in many stiffness gradient hydrogels it is not possible to know the exact stiffness to which the cells are exposed without the use of an AFM (Hetmanski et al., 2019; Lo et al., 2000; Tse and Engler, 2010). A recent report described the fabrication of easy and robust stiffness gradient hydrogels to study human adipocyte-derived stem cell behaviour (Hadden et al., 2017). However, the resulting gels are relatively thick (approx. 1 mm) and thus are not suitable for high-resolution imaging. Another report correlated diffusion of fluorescein within a PA-hydrogel mix with hydrogel stiffness, removing the need for additional AFM analyses (Koser et al., 2016); however the setup of the makeshift chamber used in this study is time consuming and is not compatible with all microscopy setups and as such limits its application and reproducibility in other labs. Nevertheless, this study demonstrated the importance of mechanical signals for axon growth, specifically the preference for axon bundles to turn towards a soft substrate.

Here, we generate thin (<100 µm) stiffness gradient hydrogels that can be easily fabricated in any laboratory, at low cost, on cell culture dishes without the need for specific equipment. These hydrogels contain fluorescently labelled beads, the density of which positively correlates with the gel’s stiffness. We generate an AFM-based correlation curve that allows researchers to assess the stiffness in every spot within the gradient of the gel simply by measuring the density of the beads using a confocal microscope. In parallel, we characterize the localization of an array of different adhesion proteins in fibroblasts and identify active α5β1-integrin as a more specific marker of fibrillar adhesions. Finally, by plating fibroblasts on physiologically relevant stiffness gradient hydrogels (0.5 – 22 kPa stiffness range) we find that fibrillar adhesions are mechano-responsive, exhibiting a logarithmic, tensin-dependent, growth in response to stiffness, rapidly increasing in length at the low stiffness regime (0.5 - 7 kPa), gradually plateauing at higher stiffness (7 – 22kPa).

## Results

### Fabrication of bead-containing stiffness gradient hydrogels

We aimed to overcome some of the limitations of currently available stiffness gradient methodologies by fabricating an easy to reproduce, low-cost and thin hydrogel suitable for high-resolution imaging. In addition, we sought a method that would allow the stiffness of the hydrogel to be measured at any given location without the need for AFM (Fig. 1). Towards this goal, we took elements from other approaches (Koser et al., 2016; Lo et al., 2000), and developed a new method to generate stiffness gradient hydrogels. We prepared two polyacrylamide (PA) solutions corresponding to the softest and the stiffest parts of our desired hydrogel gradient and included fluorescently (505/515 nm; yellow-green) labelled beads (0.1 µm carboxylated FluoSpheres) within the stiff PA solution. We then allowed the two PA mixtures to simultaneously diffuse and polymerise on a glass-bottom dish (Fig. 1A). Using this method, we consistently observed a region of bead gradient, which formed at the interface between the soft and stiff hydrogels, while other regions were either devoid of beads (corresponding to the softest hydrogel stiffness) or contained a homogenous distribution of beads (corresponding to the stiffest region of the hydrogel) (Fig. 1B, C).

**Figure 1.**
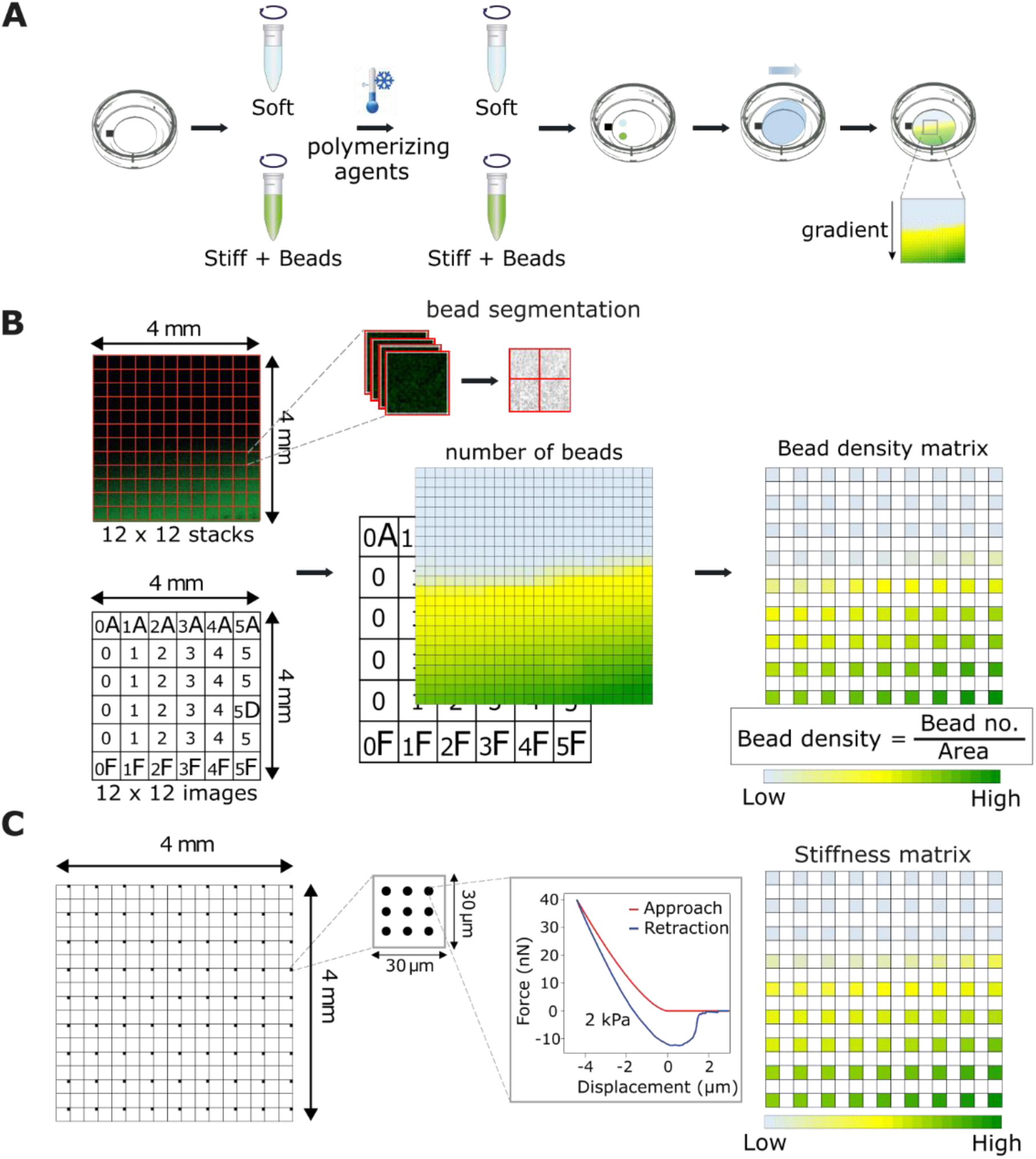
Fabrication of PA gradient hydrogels and generation of the correlation curve between fluorescent beads and stiffness. **(A)** Schematic for the fabrication of PA gradient hydrogels. A petri dish with a gridded glass-bottom well was used to fabricate the hydrogels. Two PA solutions representing the extremes of the desired hydrogel gradient were dropped onto the glass, near a pre-drawn reference mark, and allowed to mix and diffuse on the surface, leading to the formation of a gradient. The stiff PA mix also contained fluorescent beads to infer hydrogel stiffness in later steps. **(B)** A 4 mm x 4 mm region of bead gradient was selected and imaged using a spinning disk confocal microscope (12(x) x 12(y) x 7(z); total of 144 stacks). Each stack was segmented and thresholded for bead fluorescence and a 2D matrix of bead density was created. In addition, a tile scan image of the gridded glass-bottom at the same area was acquired.**(C)** Using the image of the grid, the same region of the hydrogel was located and force measurements were performed using a JPK NanoWizard® AFM system. Force measurements were carried out at different locations (0.5 mm apart in x and y coordinates when possible) within the region of interest (black squares; nine indentations distributed in 3 x 3 point grid) and the Young’s elastic modulus for each force curve was calculated. A 2D matrix with spatial distribution of stiffness was then generated. The resulting matrices from (B) and (C) were used to calculate the best fit for the correlation curve between bead density and stiffness.

### Generation of a correlation curve

We hypothesised that the concentration of beads in the hydrogel at any given point would correlate with the stiffness of the hydrogel, enabling i) rapid visual validation of the stiffness gradient with a fluorescence microscope and ii) a means to infer gel stiffness based on bead density rather than fluorescence intensity, which can be extremely variable, depending on microscope settings, and is subject to bleaching. To investigate this hypothesis, we set out to generate a correlation curve of AFM-defined stiffness versus bead density. In addition, since our protocol allows different stiffness gradients to be produced by simply changing the Young’s modulus of the two starting PA solutions, we applied our analyses to two different gradients, a wide range (2 – 60 kPa) and a narrower, softer stiffness range (0.5 – 22 kPa).

To pinpoint the same position within the hydrogel under two different imaging modalities, we prepared the hydrogels on gridded glass-bottom dishes (or used a reference mark), and then obtained a tile scan of bead distribution within the bead gradient using a spinning disk confocal microscope (Fig. 1B; see materials and methods), followed by AFM force measurements at defined points across the same area (Fig. 1C; see materials and methods). Our analyses demonstrated that in both instances AFM-defined stiffness did correlate with bead density (Fig. 2A, B). Moreover, the correlation curve for the narrower stiffness range (0.5 - 22 kPa) hydrogels could be best described as linear (Fig. 2A). In comparison, the wide-range stiffness (2 – 60 kPa) correlation curve appeared to exhibit a linear relationship between bead density and gel stiffness only in the middle ranges, whereas at the two extremes, significant changes in stiffness were not accompanied by changes in bead density, potentially reflecting the large difference between the two starting PA gels (2 kPa and 60 kPa). We found that the best model to describe this behaviour was a logit curve (Fig. 2B). Altogether, we demonstrate that it is possible to determine hydrogel stiffness based on bead density alone, bypassing the need for AFM analyses or measurements relying on fluorescence intensity.

**Figure 2.**
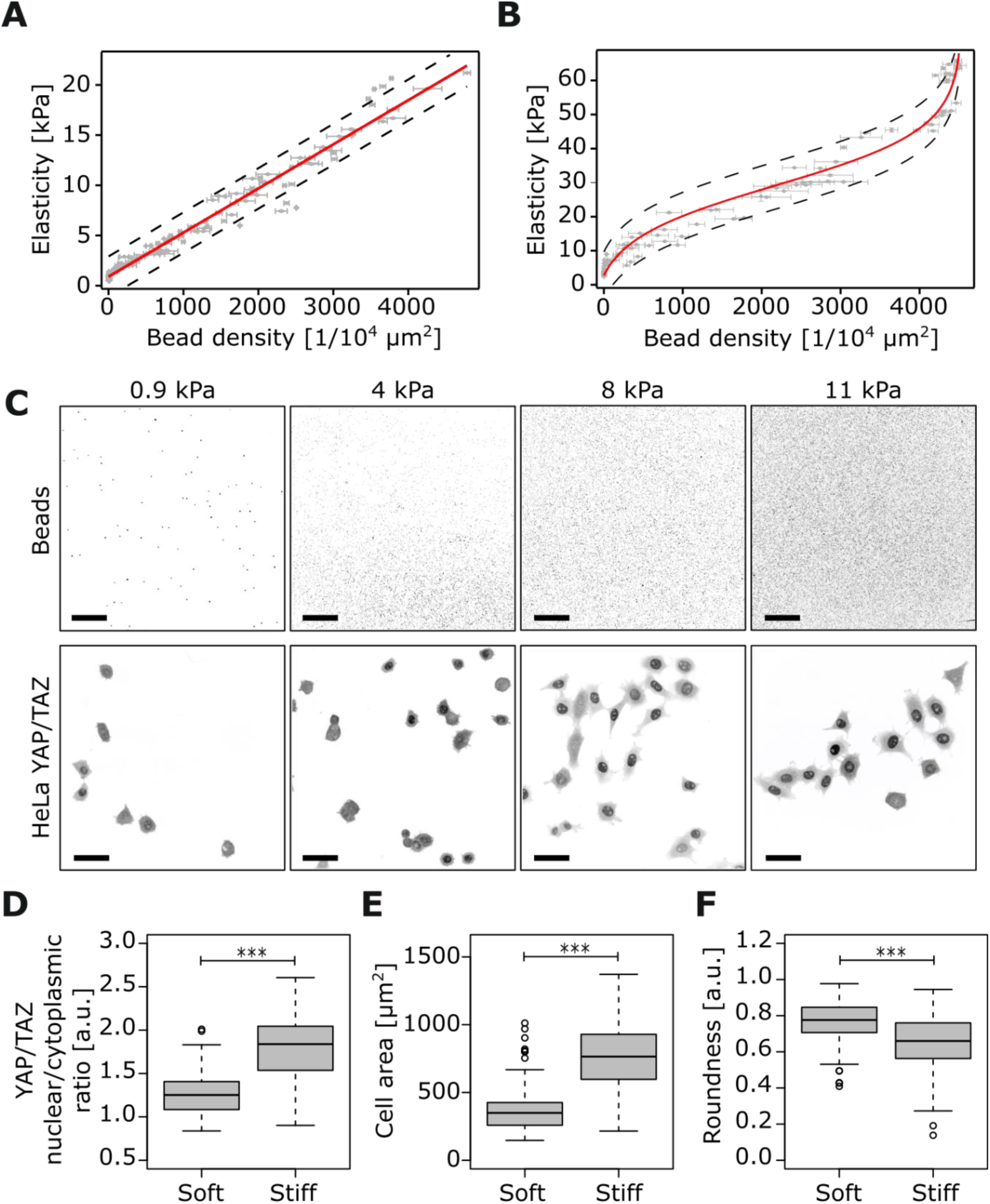
Correlation curves between bead density and stiffness and validation of hydrogel gradient. **(A)** Narrow range (0.5 kPa to 22 kPa) correlation curve. The best fit corresponds to a linear function (n = 3 independent experiments). **(B)** Wide range (2 kPa to 60 kPa) correlation curve. The best fit corresponds to a Logit function (n = 3 independent experiments). For both A and B, each data point shows the standard error (SE) for averaged bead density (horizontal error bar) and averaged stiffness (vertical error bar; nine indentation points at each location). Dashed lines at either side of the curve correspond to the 95% confidence interval (CI). **(C – F)** HeLa cells were plated on the narrow range gradient hydrogels. Representative images of the beads (first row) and YAP/TAZ intracellular localization (second row) across the hydrogel’s gradient are shown. Bead density was used to calculate the hydrogel stiffness (linear function) (C). Analysis of YAP/TAZ nuclear to cytoplasmic ratio (D), cell area (µm^2^) (E) and cell roundness (F) located on the gradient’s softer part (< 1 kPa) compared with cells located on the stiffer part (> 8 kPa) (n=2 hydrogels; 80 cells per stiff and soft part for each hydrogel; *** P < 0.001; scale bar: 50 µm).

To validate the correlation between bead density and hydrogel stiffness, we compared the real gel stiffness, measured by AFM, to the calculated stiffness, based on bead density and the best-fit calibration curve, using the wide-range stiffness gradient hydrogels. We found a high degree of correlation between AFM- and bead-density-derived stiffness measurements, i.e. AFM measurements were within the 95% confidence interval of the calibration curve (Fig. S1A).

### Biological validation of 2D-gradient hydrogels

Next, we sought to validate the biological applicability of our hydrogels by monitoring the subcellular localisation of YAP/TAZ, mechanosensitive transcriptional co-regulators, which are reported to undergo a cytoplasmic─to─nuclear switch in response to ECM stiffness (Dupont et al., 2011; Elosegui-Artola et al., 2017). Indeed, on our narrow range softer hydrogels (0.5 – 22 kPa) we observed predominately cytoplasmic YAP/TAZ localisation at the softest region of the gel measured (0.9 kPa) that became progressively more nuclear as the stiffness gradient increased (Fig. 2C). The YAP/TAZ nuclear localisation on the stiff portion of the gradient was also associated with increased cell spreading (enhanced cell area and decreased roundness) (Fig. 2D-F). These data indicate that narrow range gels could be used to monitor changes in cell morphology and to track the activation and/or subcellular localisation of different mechanosensitive proteins in response to stiffness and perhaps help identify stiffness thresholds/responsiveness in other biological processes.

### Identification of a fibrillar adhesion marker

To be able to quantitatively investigate fibrillar adhesions in respect to substrate stiffness, we set out to first identify an ideal fibrillar adhesion marker. We allowed human telomerase-immortalized fibroblasts (TIFs) to form stable adhesions on fibronectin and then determined the localisation of selected adhesion proteins, reported to be in focal or fibrillar adhesions, in a pairwise manner using a high-resolution OMX TIRF microscope and ratiometric fluorescence analysis (Zamir et al., 1999). We confirmed that tensin-1 and active α5β1-integrin (labelled with SNAKA51 antibody), previously reported to be enriched at fibrillar adhesions, demonstrate equal abundance in centrally located adhesions (Fig.3A) that are characteristic of fibrillar adhesions. These central adhesions, while rich in fibronectin (Fig 3B and Fig. S1B), were largely devoid of the focal adhesion component vinculin (Fig. S1B). Active α5β1-integrin co-localised strongly with fibronectin (Fig. 3B), whereas tensin-1 was present in prominent vinculin-positive peripheral adhesions (Fig. S1C) in addition to central adhesions, suggesting a weaker colocalization between tensin-1 and fibronectin (dual labelling with fibronectin and tensin-1 antibodies was not possible due to antibodies being raised in the same species).

**Figure 3.**
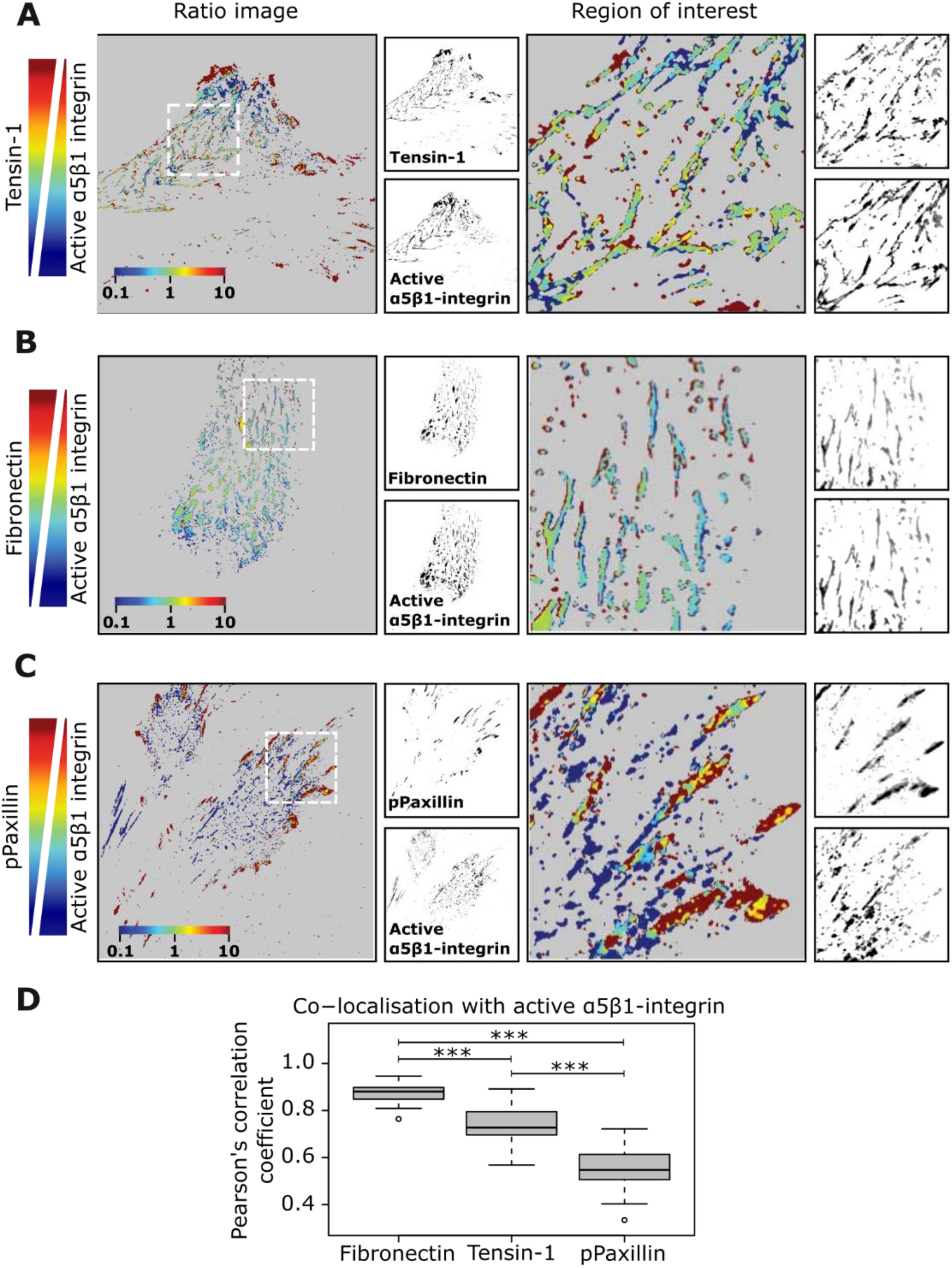
Active α5β1 integrin as a fibrillar adhesion marker. **(A – D)** TIF cells were plated on fibronectin-coated glass-bottom dishes overnight and stained for active α5β1-integrin and the indicated adhesion markers. Representative images and ratiometric analyses of colocalization between active α5β1-integrin (SNAKA51 antibody) and tensin-1 (A), active α5β1-integrin and fibronectin (B) and active α5β1-integrin and phospho-paxillin (C) and quantification of colocalization (Pearson’s coefficient) are shown (D) (n=21 to 28 cells; *** P < 0.001; ROI dimension: 20 µm x 20 µm). To obtain the Pearson’s coefficient between each pair of images, the Fiji plugin JACoP was used. The Tukey box plots display the median and the interquartile range (IQR: 25th– 75th percentile). Whiskers extend to ± 1.5 x IQR and circles represent outliers.

Intrigued by the overlap between tensin-1 and vinculin, we analyzed the distribution of tensin-1 in relation to other focal adhesion components. Dual-labelling of tensin-1 with talin confirmed that tensin-1 is co-expressed with talin in focal adhesions (Fig. S2A). In contrast, fibronectin was absent from paxillin- and talin-1-positive adhesions (Fig. S2B). Altogether our data demonstrates that tensin-1 is a component of both focal and fibrillar adhesions and thus may not be an ideal marker of fibrillar adhesions in stably adhered cells. However, active α5β1-integrin, which demonstrated a strong overlap with fibronectin in centrally located adhesions and is absent from peripheral adhesions, also showed limited colocalization with phospho-paxillin (Fig. 3C, D) and thus in line with fibrillar adhesions being viewed as phosphotyrosine poor structures within the cell (Zamir et al., 2000), may be a more appropriate fibrillar adhesion marker.

### Mechano-responsive fibrillar adhesions

To address whether fibrillar adhesions respond to rigidity, we first plated TIFs overnight on fibronectin-coated hydrogels representing two extremes of substrate stiffness (0.8 kPa, very soft; 60 kPa, very stiff). As shown previously (Yeung et al., 2005), we observed that TIFs spread more, exhibiting a flatter morphology, on the stiff versus the soft substrate (Fig.4A). We measured the length of adhesions positive for active α5β1-integrin and negative for phosphotyrosine-paxillin and found that on a soft substrate fibroblasts have small, often dot-like adhesions, whereas on a stiff substrate the adhesions are primarily longer resembling more typical fibrillar adhesions (Fig. 4A, B). Next, we plated cells on the narrower stiffness gradient hydrogels (0.5 – 22kPa) and monitored fibrillar adhesion formation. We made the interesting observation that the length of active α5β1 integrin adhesions positively correlates with the stiffness of the substrate (Fig. 4C, D). This increase in adhesion length could be best described with a logarithmic distribution - rapid increase at lower stiffness (1-7 kPa), followed by a more gradual increase and finally plateau at higher stiffness (7-22 kPa) reaching a maximum length of approx. 3.5 µm in our system.

**Figure 4.**
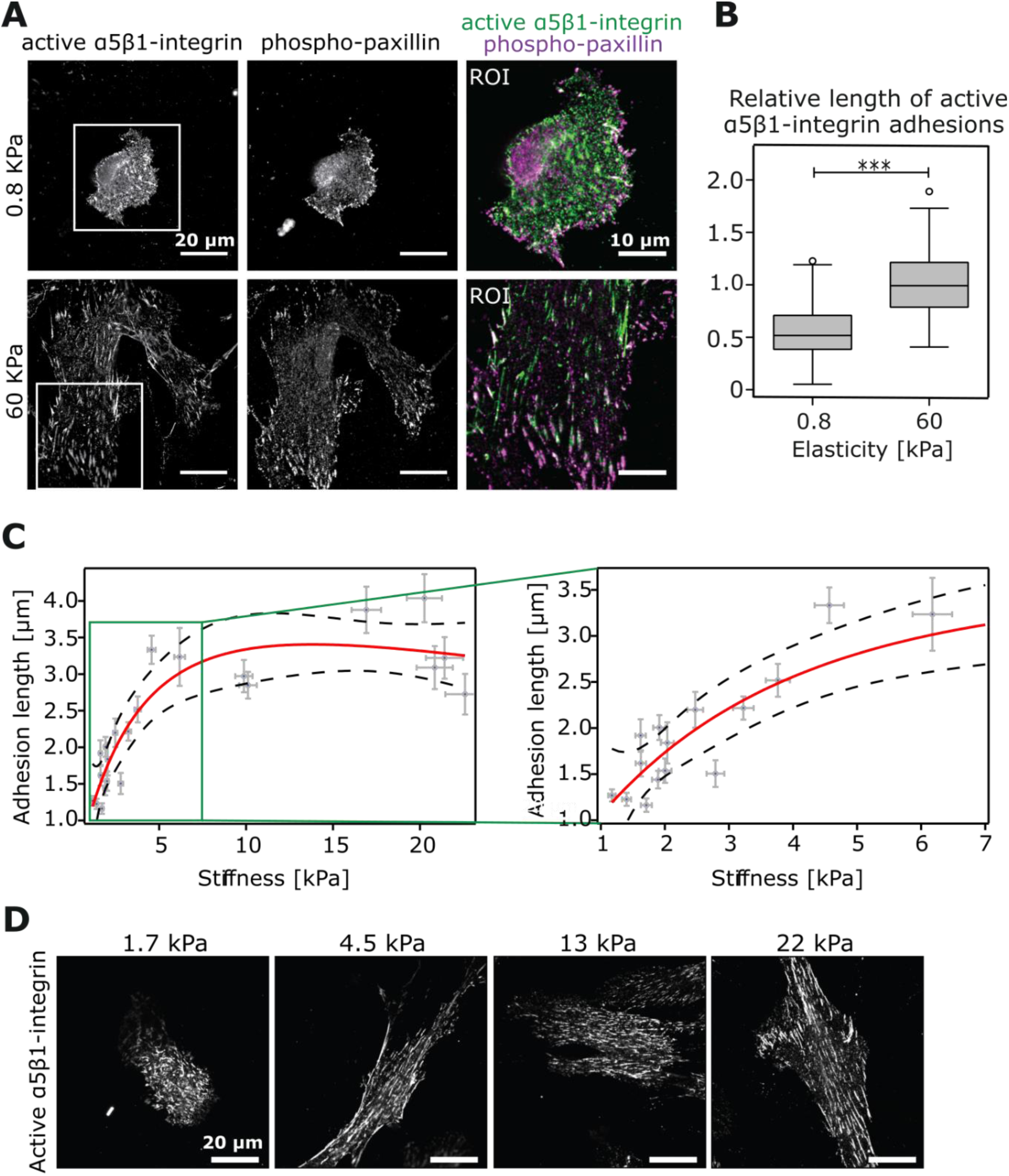
Active α5β1 integrin adhesions respond to changes in stiffness. **(A)** Representative images of TIFs plated on uniform stiffness hydrogels (0.8 kPa or 60 kPa) and stained for active α5β1-integrin and phospho-paxillin (scale bar: 20 µm). **(B)** Tukey box plot of α5β1-integrin adhesion length in µm from A (n=2 independent experiments, 9-10 cells analysed per experiment; 199 adhesions (0.8 kPa) and 211 adhesions (60 kPa); *** P < 0.001). **(C)** Correlation analysis between stiffness (kPa) and α5β1-integrin adhesion length (µm) in TIFs plated on a stiffness gradient hydrogel (0.5 kPa to 22 kPa) (each data point represents mean adhesion length ± SD analysed from one field of view). Red line represents the line of best fit, and the dashed lines at either side correspond to the 95% CI. Green rectangle shows the region of interest highlighted in the right-hand-side panel. **(D)** Representative images of TIFs stained for active α5β1 integrin (SNAKA51 antibody) from (C) across the stiffness gradient (scale bar: 20 µm).

### Tensins support stiffness-induced adhesion elongation

Tensins bind directly to the integrin β1-tail and stabilize integrins on the cell surface (Lo et al., 1994; Torgler et al., 2004). In addition, tensins maintain β1-integrin activity in fibrillar adhesions following initial talin-dependent switching of the receptor into an active conformer (Georgiadou et al., 2017). Moreover, a reduction in fibrillar adhesion number, in tensin-depleted cells or upon AMPK activation, correlates with significantly reduced traction-forces on fibronectin (Georgiadou et al., 2017), indicating that fibrillar adhesions transduce forces to the ECM. To test whether tensins are required for the stiffness-dependent increase in active α5β1 integrin adhesion length, we silenced tensin-1 expression using siRNA oligos that we have previously validated for specificity with rescue experiments (Georgiadou et al., 2017). Interestingly, tensin silencing, validated with qRT-PCR (Fig 5A), clearly reduced active α5β1 integrin adhesion length in cells plated on the stiffness-gradient gels when compared to the control-silenced cells (Fig 5B, C). These data demonstrate that while tensins may not be restricted to fibrillar adhesions, they are important for active α5β1 integrin adhesion elongation on a range of matrix rigidities.

**Figure 5.**
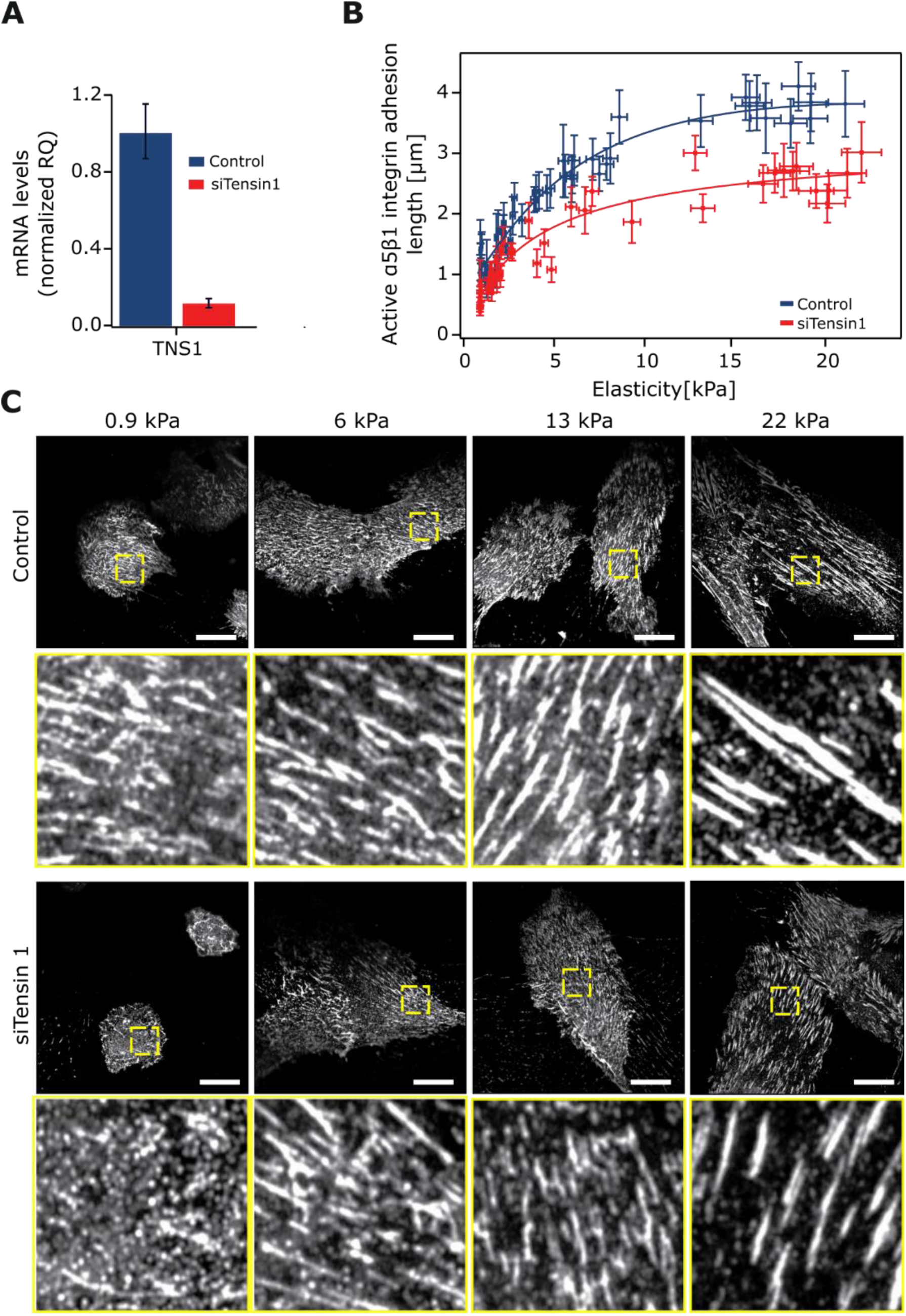
Tensins are required for adhesion elongation in response to stiffness. **(A)** Taqman qPCR analysis of tensin-1 silencing efficiency in TIF cells. **(B – C)** siTensin-1 and control-silenced TIFs were plated on stiffness gradient hydrogels and stained for active α5β1 integrin. Comparison of active α5β1-integrin adhesion length across a hydrogel stiffness gradient between Tensin-1-silenced TIFs (n=3) and controls (n=2) (B; data are mean ± SD) and example images (C) are shown. Yellow insets denote magnified regions of interest of α5β1-integrin adhesions (Scale bar: 20 µm).

## Discussion

Many fundamental cellular processes including proliferation, migration and differentiation are precisely regulated by cues emanating from a dynamic and heterogeneous microenvironment. These cues include fluctuations in the biochemical composition as well as the biophysical properties (viscoelasticity, topography and architecture) of the surrounding ECM.

Several methods have been developed to reduce the complexity of the microenvironment in order to imitate the effect of ECM stiffness on cell behaviour (Hadden et al., 2017; Koser et al., 2016; Lo et al., 2000; Tse and Engler, 2010; Vincent et al., 2013) and primarily involve the production of PA hydrogel-based stiffness gradients. While all of these techniques have their own unique advantages, limitations include production of thick hydrogels that are not compatible with high-resolution imaging or for live-cell dynamics, the need for AFM verification of stiffness for individual experiments and the use of fine-tuned techniques for specific stiffness gradients that reduce reproducibility in other laboratories. Here, we set out to develop a simple and cost-effective method to produce stiffness gradients based on previous approaches (Koser et al., 2016; Lo et al., 2000). We demonstrate that our technique is flexible and can be applied to produce different rigidity gradients without the need for specialised equipment. The resulting hydrogels can be used as a reductionist model to image and dissect mechanosensitive pathways and signalling in cells. We show that within a 0.5 – 22 kPa range, cell spreading increases and YAP/TAZ localisation becomes progressively nuclear with increasing stiffness. While these results are expected, these proof-of-concept data indicate that our microscopy amenable hydrogels could potentially be used to simultaneously chart the effect of substrate stiffness heterogeneity on two or more proteins within the same dish or to track dynamic changes in individual cells when they encounter different mechanical cues. Whether precise stiffness thresholds, for example for inducing complete YAP/TAZ nuclear translocation, could be determined for different cell lines, remains to be investigated but would be fundamental in our understanding of mechanosignalling in development and disease.

We also used our stiffness gradient hydrogels to explore whether fibrillar adhesions, the main sites of fibronectin fibrillogenesis, respond to changes in ECM rigidity. We demonstrate that in TIFs, fibrillar adhesion length, identified by active α5β1 integrin staining, increases rapidly up until approximately 7 kPa. After this point, adhesion lengthening decelerates and eventually becomes relatively stable, suggesting that fibrillar adhesions are indeed mechanosensitive. Importantly, we find this mechanosensitive adhesion lengthening to be tensin-dependent. Recently, tensins have been implicated in supporting integrin activity and traction-forces in fibroblasts in vitro (Georgiadou et al., 2017) in addition to integrin activity in vivo in the myotendinous junctions of drosophila flight muscles (Green et al., 2018). The exact nature of how fibrillar adhesions retain their connection to the actin cytoskeleton, perhaps through integrin─tensin interaction, remains to be investigated. However, our data show that these structures respond to gradual changes in ECM rigidity.

In our set-up, we opted to use bead density rather than fluorescence intensity as a readout of hydrogel stiffness. We show that while there is a linear correlation between bead density and hydrogel stiffness at narrower stiffness gradients (0.5 – 22 kPa), at wider stiffness gradients a logit fit appears to be a more accurate representation of the relationship (2 – 60 kPa). This is an important consideration that has not been highlighted previously, for example, when fluorescein was used as a means to measure hydrogel stiffness (range of 0.1 – 10 kPa; (Koser et al., 2016)). In addition, we believe that the substitution of fluorescein intensity with the analysis of bead density (our method) to measure stiffness, is a more flexible and viable approach, as we are not relying on fluorescence intensity, which as a read-out can be highly variable depending on bleaching rate and on the imaging modality used.

Fibronectin structure and function undergo mechano-regulated alterations (Craig et al., 2001; Smith et al., 2007) that could for example influence fibronectin-dependent assembly of other ECM components such as collagen (McDonald et al., 1982; Saunders and Schwarzbauer, 2019; Velling et al., 2002). However, the notion that, through mechanosensitive fibrillar adhesions, fibronectin remodelling may also be subject to regulation by substrate rigidity has received less attention. The stiffness-dependent lengthening of fibrillar adhesions, observed here, has potentially important implications in tissue fibrosis (Chen et al., 2014; Pelouch et al., 1993), cancer (Cox and Erler, 2011) and drug resistance. In the context of cancer, this process may impinge on fibronectin-guided invasion of cancer cells in the tumour microenvironment (Oudin et al., 2016) or on nutrient sensing through the modulation of integrin α5β1 endocytosis and recycling (Georgiadou and Ivaska, 2017; Rainero et al., 2015).

## Acknowledgements

We thank P. Laasola and J. Siivonen for technical assistance and the Cell Imaging and Cytometry core facility at Turku Bioscience Centre, University of Turku for help with imaging. This study has been supported by the University of Turku Doctoral Programme in Molecular Life Sciences (DPMLS) (A.I.), the Academy of Finland (M.G., J.I.), the Sigrid Juselius Foundation (J.I.), the Cancer Society of Finland (J.I.) and by an ERC consolidator grant (AdheSwitches, 615258; J.I.).

## Author contributions

Conceptualization: N.B., M.G. and J.I; Methodology: N.B., M.G., C.G., A.I., and J.I; Investigation: N.B., M.G., C.G.; resources: J.I; Writing original draft: M.G., H.H; writing – reviewing: N.B., M.G., C.G., A.I. and J.I; Visualization: N.B., M.G., H.H; Supervision: J.I; Funding acquisition: J.I.

## Supplementary Figures

**Supplementary figure 1.**
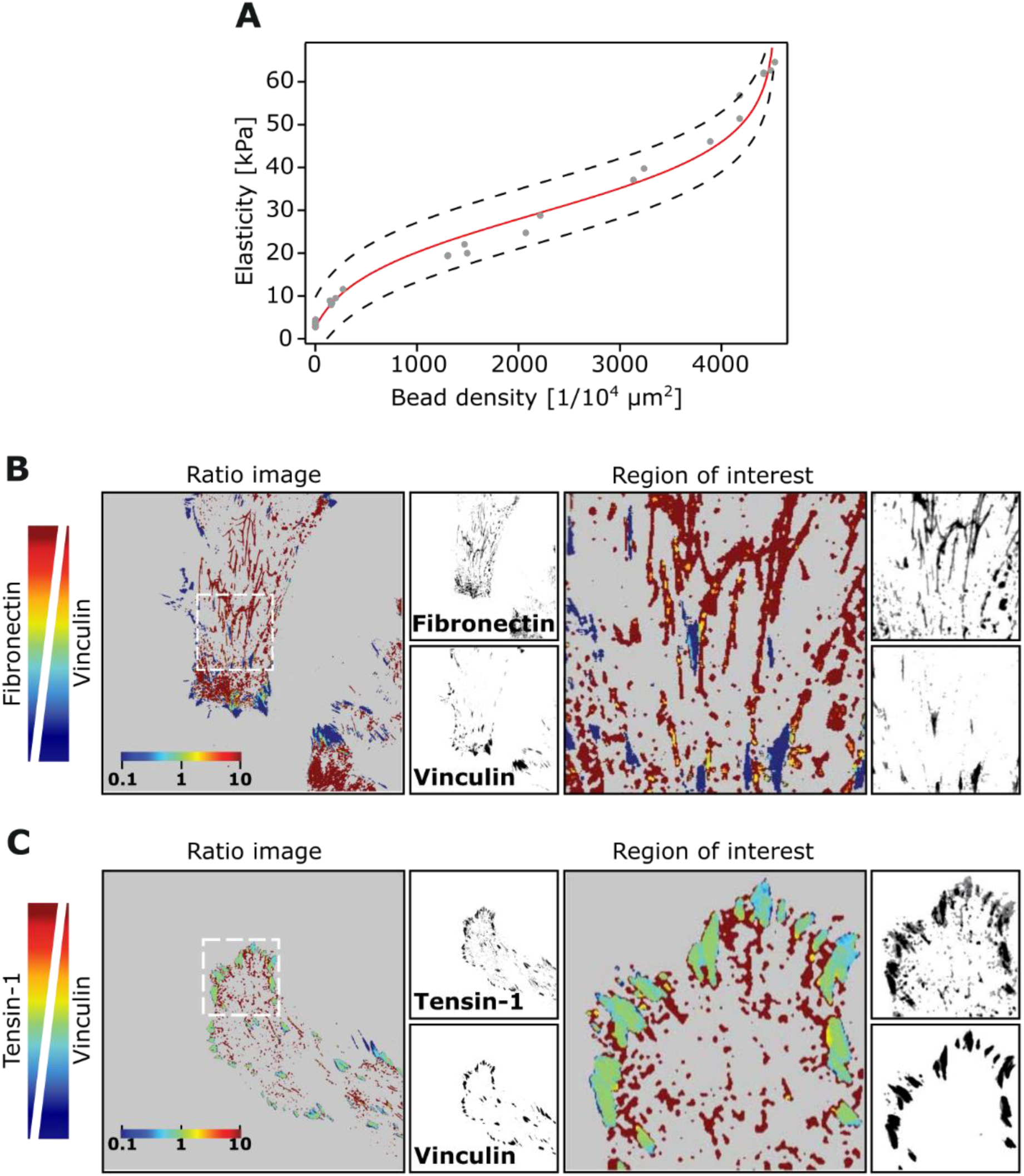
**(A)** AFM validation of calibration curve-derived stiffness values. AFM was used to measure hydrogel stiffness at different regions within the stiffness gradient hydrogel and the results were compared to the values derived from the calibration curve. Red line represents the calibration curve, grey circles are the AFM measurements at the indicated bead density and the dashed lines at either side of the curve correspond to the 95% CI. **(B – C)** TIF cells were plated on fibronectin-coated glass-bottom dishes overnight and stained for the indicated adhesion markers. Representative images and ratiometric analyses of colocalization between vinculin and fibronectin (B) and vinculin and tensin-1 (C) are shown.

**Supplementary figure 2.**
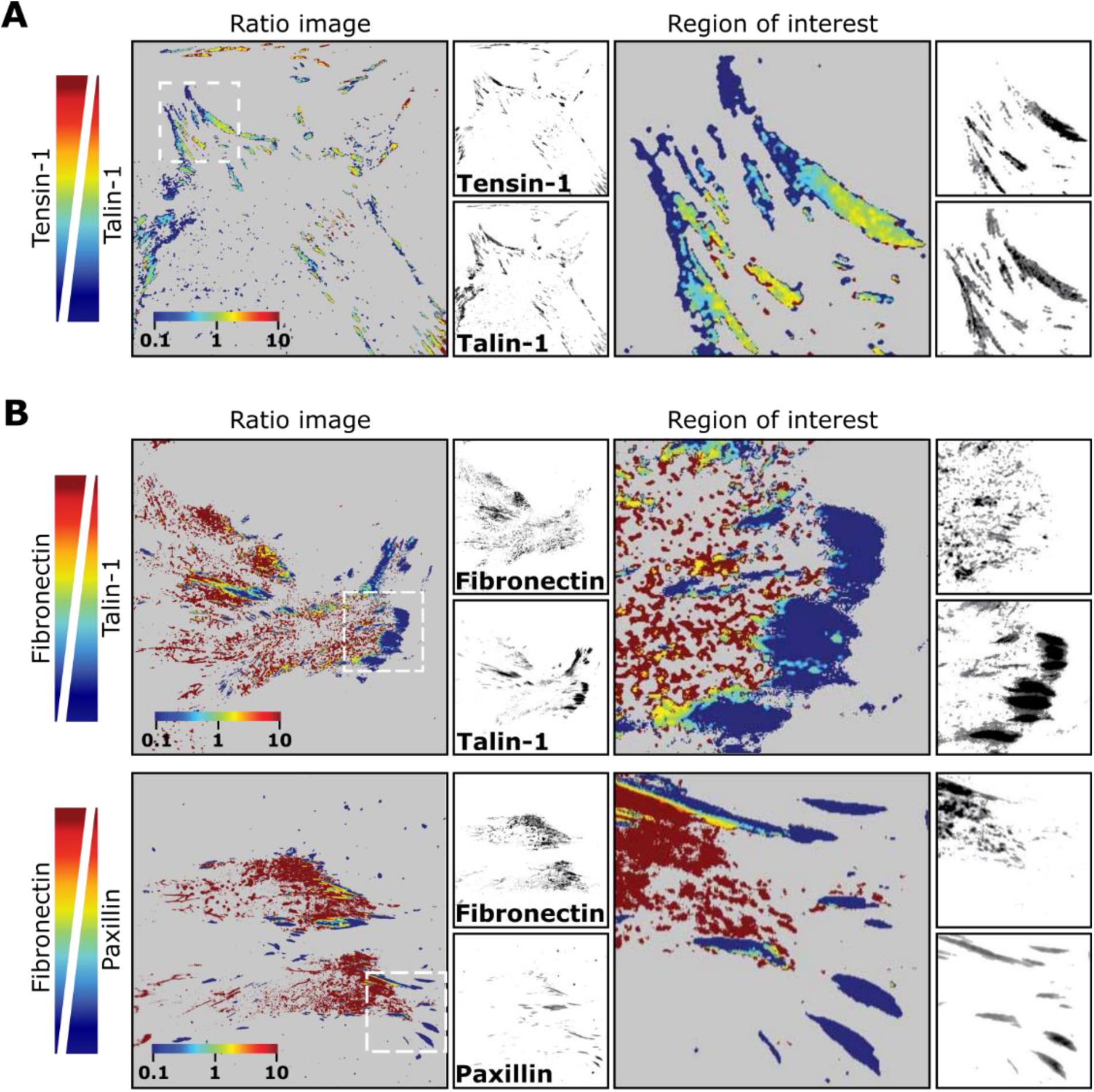
**(A – B)** TIF cells were plated on fibronectin-coated glass-bottom dishes overnight and stained for the indicated adhesion markers. Representative images and ratiometric analyses of colocalization between tensin-1 and talin-1 (A) and between fibronectin and talin-1 or paxillin (B) are shown.

## Materials and methods

### Fabrication of PA gradient hydrogels

Glass-bottom dishes (0.13 - 0.16 thickness; 14 mm diameter, Cellvis, D35-14-1-N) were treated for 20 min at room temperature (RT) with 200 µl of Bind-silane solution—a mixture of 714 µl 3-(Trimethoxysilyl)propyl methacrylate (3-TMP, Sigma-Aldrich, M6514), 714 µl of acetic acid in 10 ml of 96% ethanol. This mix was used to covalently attach PA hydrogels to the glass surface and to prevent hydrogel detachment. After the Bind-silane was aspirated, the glass surface was washed twice with ethanol and left to dry completely. A reference mark was also manually drawn on the bottom of the dish (Fig. 1A).

Two PA pre-mix solutions, one soft (0.5 kPa or 2 kPa) and one stiff (20 kPa or 60 kPa), were prepared to create rigidity gradients of ~ 0.5 – 20 kPa and ~ 2 ─ 60 kPa. The desired Young’s modulus (*E*) of the pre-mixes was adjusted by mixing pre-defined ratios of 40% (w/v) acrylamide monomer (Sigma-Aldrich, A4058) and 2% (w/v) N, N methyl-bis-acrylamide cross-linker (Sigma-Aldrich, M1533) in PBS (Table 1). A standard volume (1.7 µl, 3.6 × 10^10^ beads/µl) of fluorescently labelled (505/515 nm) beads (0.1 µm carboxylated FluoSpheres; ThermoFisher, F8803) was sonicated (3 min) and added into the stiff pre-mix. Both PA pre-mix solutions, soft and stiff, were briefly vortexed and kept on ice to avoid fast polymerization in later steps.

**Table 1.**
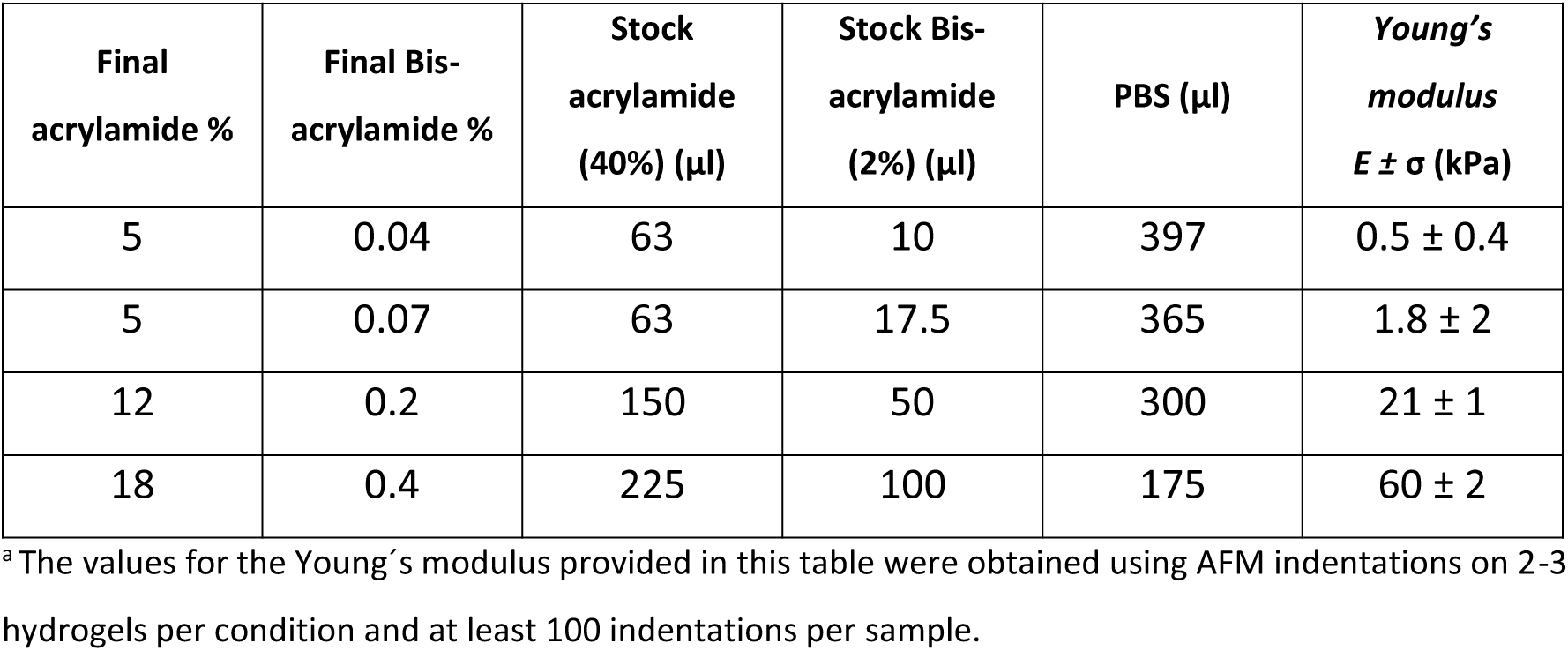
Relative acrylamide and bis-acrylamide concentrations for the fabrication of uniform (constant *E*) hydrogels and expected Young’s modulus after polymerization ^a^

| Final acrylamide % | Final Bis-acrylamide % | Stock acrylamide (40%) (μl) | Stock Bis-acrylamide (2%) (μl) | PBS (μl) | Young's modulus $E \pm \sigma$ (kPa) |
| --- | --- | --- | --- | --- | --- |
| 5 | 0.04 | 63 | 10 | 397 | $0.5 \pm 0.4$ |
| 5 | 0.07 | 63 | 17.5 | 365 | $1.8 \pm 2$ |
| 12 | 0.2 | 150 | 50 | 300 | $21 \pm 1$ |
| 18 | 0.4 | 225 | 100 | 175 | $60 \pm 2$ |

<sup>a</sup> The values for the Young's modulus provided in this table were obtained using AFM indentations on 2-3 hydrogels per condition and at least 100 indentations per sample.

Polymerization of the soft pre-mix was started immediately after the previous step by addition of 5 µl 10% ammonium persulphate (APS; BioRad) and 1 µl N, N, N’, N’-tetramenthylethylenediamine (TEMED; Sigma T-9281) to the solution. The polymerizing soft mixture was quickly vortexed and a 7.8 µl droplet of the solution was pipetted on top of the glass-bottom well approximately 3 mm across and 1 mm above the reference mark. The same polymerisation procedure was repeated with the stiff pre-mix and a 7.8 µl droplet of the solution was placed approximately 2 mm below the soft PA droplet. A circular coverslip (13 mm) was then placed on top of the droplets by gently dropping it from the reference mark’s edge towards the opposite side of the glass well, leading to in situ mixing of PA gels and diffusion across the dish.

The hydrogel was left to polymerize for 1 h at RT. Upon polymerization the gel was covered with PBS for 5 min before the coverslip was carefully removed with a bent needle. Lastly, the hydrogel was washed with PBS to remove any remaining unpolymerized PA, and then immersed in PBS and stored at 4°C until needed. Based on the employed volume of PA and the dimension of the glass-bottom dishes, we could estimate a hydrogel thickness of approximately 100 µm, confirmed using a spinning disk microscope (data not shown).

### Generation of stiffness gradient correlation curves from PA hydrogels loaded with fluorescent beads

Correlation curves were generated for a wide stiffness range hydrogel (2 – 60 kPa) and a narrow stiffness range hydrogel (0.5 – 22 kPa). For this purpose, hydrogels were prepared on gridded glass-bottom dishes (Cellvis, D35-14-1.5GO) as above to allow the same area to be located under different microscopes (SDC and AFM).

#### Analysis of bead number

The bead gradient within the hydrogel was pinpointed using a spinning disk confocal microscope (3i CSU-W1) equipped with a 40X objective lens (C-Apochromat 40X/1.1 NA; Zeisss) and a sCMOS (Hamamatsu Orca Flash 4; Hamamatsu Photonics) camera. A tile scan (12(x) x 12(y) x 7(z) images) covering an area of 4 mm x 4 mm was acquired (488 nm laser line, intensity: 800 W/cm^2^; GFP 510-540 nm emission filter). The z-upper-limit for each stack was set 1 µm underneath the gel’s surface resulting in 144 stacks of 324.48 µm X 324.48 µm X 10 µm in size. The focal plane of the microscope was then changed to focus on the gridded glass-bottom, and a tile scan of bright-field images (12(x) x12(y)) covering the same region as the beads was acquired (Fig 1B).

A semi-automatic Fiji macro with custom scripts was then used to process the acquired images. Briefly, for each stack a maximum intensity projection was produced and then segmented, with the appropriate threshold, into a 2 x 2 grid (total 576 images from the original 144 stacks), allowing a more accurate quantification of the beads within the same image. A custom Python script was then used to calculate the density of beads per area unit (1/10^4^ µm^2^) and to create a 2D matrix displaying the spatial distribution of bead density (Fig 1B).

#### AFM analysis of hydrogel stiffness gradient

Hydrogel elasticity was assessed using a JPK NanoWizard with a CellHesion module mounted on a Carl Zeiss confocal microscope, Zeiss LSM510 (AFM; JPK instruments) and silicon nitride cantilevers (spring constant: 1Nm^−1^, spherical 10 µm diameter tip; Novascan Technologies). The cantilever spring constant and deflection sensitivity were calibrated in fluid via the thermal noise method (Hutter & Bechhoefer, 1993). Prior to distance force measurements, a CCD camera mounted on the AFM was used to visualize the grid of the culture dish and to locate the same 4 mm x 4 mm region of interest previously imaged with the spinning disk microscope. Force measurements were then carried out at different locations (0.5 mm apart in x and y coordinates) within the region of interest. In each location, nine indentations distributed in a 3 x 3 point grid (30 µm x 30 µm) were performed. The elastic modulus for each force curve was calculated using JPK data processing software (JPK DP version 4.2) assuming a Hertz model of impact (Fig 1C).

A custom Python script was then used to consolidate all elasticity measurements from multiple files into a single file, to calculate the mean between the nine stiffness values obtained per location and to create a 2D matrix displaying the spatial distribution of stiffness (Fig 1C).

#### Correlation between bead density and AFM-defined hydrogel elasticity

To assess the correlation between bead density and hydrogel elasticity, the tile scan of the grid was overlaid with the bead density matrix. By doing this, it was possible to identify the bead location corresponding to the point where the elasticity measurements were taken. The Igor Pro software (IgorPro 6.37, Wavemetrics) was then used to plot bead density against elasticity and to calculate the best fitting curve for the data. In both cases, wide range (2 to 60 kPa) and narrow range (0.5 to 22 kPa) gradients, data from three independent hydrogels was processed as previously described and combined to generate the two final correlation curves.

The best fit for the narrow range correlation curve (0.5 – 22 kPa) corresponded to the following linear function:

- *y* = *a* ∗ *x* + *b*
- *y* = 0.0044 ∗ *x* + 0.903

where *y* corresponds to stiffness, *x* to bead density (number of beads in an area of 100 µm x 100 µm), and the fitted constants *a* and *b* to the slope and the intercept respectively.

The best fit for the wide range correlation curve (2 – 60 kPa) corresponded to the following Logit function:

- 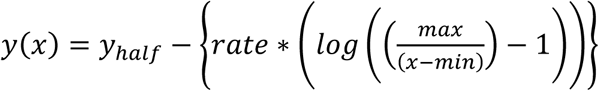
- 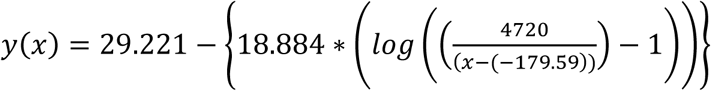

where *y* corresponds to the elastic modulus, *x* to bead density (number of beads in an area of 100 µm x 100 µm), *y*_*half*_ (fitted constant; 29.221 ± 4.67E-15) is the half point of the logit where there is a change in curvature, *rate* (fitted constant; 18.884 ± 6.19E-15) is the rate at which the elastic modulus is increasing and *min* and *max* (fitted constants; 4720 ± 0.00 and −179.59 ± 1.66E-13, respectively) correspond to the limit conditions in the abscissa. These equations were used in ensuing experiments to calculate hydrogel elasticity.

### Hydrogel activation and functionalization

For functionalization, 0.2 mg/ml Sulfo-SANPAH (Thermo Scientific, 22589) and 2 mg/ml N-(3-Dimethylaminopropyl)-N′-ethylcarbodiimide hydrochloride (EDC) (Sigma, 03450) in 50 mM HEPES were added on top of the hydrogels and incubated for 30 min at RT, protected from light, and with gentle agitation. Gels were then placed into a UV-chamber for 10 min to activate the Sulfo-SANPAH and finally washed three times with PBS prior to overnight coating with the indicated ECM molecule/s at 4°C.

### Cell culture

Human cervical adenocarcinoma (HeLa) cells were maintained in high glucose Dulbecco’s Modified Eagle Medium (DMEM) supplemented with 10% Fetal Bovine Serum (FBS), 1% non-essential amino acids, 1% 2 mM L-glutamine and 1% Penicillin-Streptomycin (Pen-Strep). All cells were routinely tested for mycoplasma contamination. Human telomerase-immortalized fibroblasts (TIFs) were a gift from Jim Norman (Beatson Institute, Glasgow, UK) and were cultured in DMEM 4500 supplemented with 20 % FBS, 2 mM L-glutamine and 20 mM Hepes buffer (Sigma-Aldrich).

### siRNA transfections

siRNA silencing was performed using 20 nM siRNA oligos and Lipofectamine® RNAiMAX Reagent (ThermoFisher Scientific) according to manufacturer’s protocol and cells were cultured for 3 days before the experiments. The siRNA against human tensin1 (L-009976, smartpool) and the control siRNA (D-001810-10, non-targeting Pool) were ON-TARGETplus siRNAs from Dharmacon, ThermoFisher Scientific.

### Antibodies, compounds and reagents

The following antibodies were used: mouse anti-YAP/TAZ (sc-101199), anti-tensin1 (SAB4200283, Sigma-Aldrich), anti-fibronectin (F3648, Sigma-Aldrich), anti-vinculin (clone hVIN-1, V9131), anti-talin (clone 8d4, T3287; Sigma-Aldrich), anti-paxillin (612405; BD Biosciences), anti-phosphotyrosine (Y118)-paxillin (Cell Signaling) and anti-GAPDH (Mab6C5, 5G4, HyTest). Anti-human active α5-integrin (SNAKA51) was a gift from Martin Humphries, University of Manchester, UK (Clark et al., 2005). Phalloidin Atto 647N (65906) was obtained from Sigma-Aldrich. AlexaFluor®-conjugated secondary antibodies (488, 555, 568, 647-conjugated anti-mouse, rabbit and rat antibodies, ThermoFisher Scientific) were used in immunofluorescence and flow cytometry. Bovine plasma fibronectin (FN; 341631) was purchased from Merck, Millipore and collagen type I (from calf skin) from Sigma-Aldrich (C8919).

### YAP/TAZ immunofluorescence

HeLa cells were seeded (4 × 10^3^ cells) for 24 h on gradient hydrogels pre-coated (overnight, 4°C) with 2.5 µg/ml fibronectin + 2.5 µg/ml collagen I. Cells were fixed with a final concentration of warm 4 % PFA added straight into the media at RT. Blocking and permeabilization was performed with 0.3% Triton-X in 10% horse serum for 15 min. After washing with PBS, cells were incubated overnight at 4 °C with the indicated primary antibody, mouse anti-YAP/TAZ (1:100), in 10% horse serum. Cells were then washed with PBS and incubated with AlexaFuor 568-conjugated secondary antibody (1:200; 1.5 h at RT), Phalloidin Atto 647 (1:200) and DAPI (1:1000). Finally, cells were washed and kept in PBS until imaging.

HeLa cells were imaged on a 3i (Intelligent Imaging Innovations) Marianas spinning disk confocal microscope (Zeiss), with a C-Apochromat 40X/1.1 numerical aperture (NA) water immersion objective (Zeiss) and sCMOS Orca camera (Hamamatsu Photonics). A semi-automatic custom macro script in ImageJ (Fiji) was used to determine YAP/TAZ nuclear to cytoplasmic intensity ratio. Briefly, maximum intensity projections were created and the nucleus (defined by DAPI staining) and cytoplasm (region corresponding to a 1 µm ring around the nucleus, excluding DAPI staining) were segmented by drawing one line around the DAPI staining (nucleus) and another line 1 µm away apart from DAPI staining. YAP/TAZ mean intensities were then calculated in the different regions. Cell area was calculated from maximum intensity projections of actin staining in ImageJ. Hydrogel stiffness was determined as described above using bead density and the linear equation (*Y=0.0044x (x) + 0.903)*.

### Ratiometric analysis of adhesions pairs in TIFs

TIF cells were seeded overnight on glass-bottom dishes (MatTek Corporation) pre-coated with 10 µg/ml fibronectin (overnight at 4°C), fixed and permeabilized with 4% PFA and 0.2 % Triton-X for 10 min, blocked with 1 M Glycine for 30 min, washed and then incubated with the indicated primary antibodies for another 45 min. Following further washes, cells were incubated with Alexa-conjugated secondary antibodies (6 μg/ml), Phalloidin–Atto 647N (1:200) and 0.5 μg/ml DAPI in PBS for 30 min. Finally, cells were washed with PBS and Milli-Q water and imaged with the indicated microscope.

Ratiometric analysis was performed using a modified version of a previously described protocol (Zamir et al., 1999). In short, two-colour images of TIFs stained with the proteins of interest were first processed to remove background and noise. Using the “subtract background” and the “threshold” functions of ImageJ software (NIH) a mask was created setting to zero all pixels below threshold and maintain the values of pixels above threshold. For accuracy, each of the labelled channels was processed separately. Ratio images were then calculated using the open source software R (R Core Team) and by simply diving the values pixel by pixel. Given that there exist multiple pixels with a zero value in both channels/labels, we defined a multiple case scenario to calculate the ratio image: 1) A resulting value of zero was assigned whenever the pixel in both channels/labels was zero. 2) A value of 0.1 was assigned whenever the ratio between the pixel in label A (numerator) and the pixel in label B (denominator) was ≤ 0.1. 3) A value of 10 was assigned whenever the ratio between the pixel in label A (numerator) and the pixel in label B (denominator) was ≥ 10, or in the case the numerator was >0 and the denominator was zero. 4) In all the remaining cases the pixel was assigned the ratio value between the numerator and the denominator pixel. After all ratio values were calculated and assigned, the images were displayed in log scale using a colour look-up table (Jet2 for all pixels >0 and grey for pixel values of 0), such representation allows to present ratio value variations over two orders of magnitude (from 0.1 to 10).

### Analysis of adhesion length

TIF cells were plated overnight on hydrogels (either 0.8 kPa, 60 kPa or 0.5 – 20 kPa stiffness gradient hydrogels) precoated with 10 µg/ml fibronectin and stained for active α5β1 integrin (SNAKA51 antibody) as described. The laser scanning confocal microscope (CLSM, Zeiss LSM 880 AiryScan) with LD LCI Plan-apochromat 40X/1.2 (NA) and super-resolution AiryScan detector was used to image fibrillar adhesions in cells at different locations across the bead gradient. Adhesion length was then manually measured in Fiji by using the freehand measuring tool. The mean adhesion length and standard deviation was calculated for each cell.

### Statistical analysis

The Student *t-*test (two-tailed equal variance) was used for statistical analysis.

